# Cryopreservation of cerebrospinal fluid cells preserves transcriptomics integrity for single-cell analysis

**DOI:** 10.1101/2023.08.10.552735

**Authors:** Mahesh Kodali, Jerry Antone, Eric Alsop, Rojashree Jayakumar, Khushi Parikh, Paula Sanchez-Molina, Bahareh Ajami, Steven Arnold, Kendall Jensen, Sudeshna Das, Marc S. Weinberg

## Abstract

Cerebrospinal fluid (CSF) matrix biomarkers have become increasingly valuable surrogate markers of neuropsychiatric diseases in research and clinical practice. In contrast, CSF cells have been rarely investigated due to their relative scarcity and fragility, and lack of common collection and cryopreservation protocols, with limited exceptions for neurooncology and primary immune-based diseases like multiple sclerosis. the advent of a microfluidics-based multi-omics approaches to studying individual cells has allowed for the study of cellular phenotyping, intracellular dynamics, and intercellular relationships that provide multidimensionality unable to be obtained through acellular fluid-phase analyses. challenges to cell-based research include site-to-site differences in handling, storage, and thawing methods, which can lead to inaccuracy and inter-assay variability. In the present study, we performed single-cell RNA sequencing (10x Genomics) on fresh or previously cryopreserved human CSF samples from three alternative cryopreservation methods: Fetal Bovine Serum with Dimethyl sulfoxide (FBS/DMSO), FBS/DMSO after a DNase step (a step often included in epigenetic studies), and cryopreservation using commercially available Recovery© media. In comparing relative differences between fresh and cryopreserved samples, we found little effect of the cryopreservation method on being able to resolve donor-linked cell type proportions, markers of cellular stress, and overall gene expression at the single-cell level, whereas donor-specific differences were readily discernable. We further demonstrate the compatibility of fresh and cryopreserved CSF immune cell sequencing using biologically relevant sexually dimorphic gene expression differences by donor. Our findings support the utility and interchangeability of FBS/DMSO and Recovery cryopreservation with fresh sample analysis, providing a methodological grounding that will enable researchers to further expand our understanding of the CSF immune cell contributions to neurological and psychiatric disease.

## INTRODUCTION

The lumbar puncture (LP) is a safe ambulatory procedure for collecting cerebrospinal fluid (CSF) [1]. LPs are classically applied to diagnosis of neurological diseases such as communicating hydrocephalus, autoimmune or infectious encephalitis, central nervous system vasculitis, and cancer [2]. Improvements in biomarker characterization [3] and their ultrasensitive detectability have increased the utility and promise of CSF for aiding in diagnosis for neuropsychiatric disorders [4], and for neurodegenerative diseases like Alzheimer’s disease (Aβ42, t-Tau, and p-Tau181) [5], motor neuron disorders [6], and prion disease [7]. Given the recent United States Food and Drug Administration approval of several monoclonal antibody-based treatments against amyloid-β in Alzheimer’s disease, and the requirement of either CSF- or amyloid-PET imaging-based biomarker thresholds for receiving lecanemab [8] and likely other similar forthcoming treatments, we anticipate that the clinical utilization of the lumbar puncture is likely to increase in the United States in coming years.

CSF is a clear fluid produced by the choroid plexus and ependymal cells lining the brain’s ventricles. It is in continuous interchange with the brain’s interstitial fluid as it circulates through subarachnoid and glymphatic spaces [9]. The acellular and cellular composition of CSF differs greatly from that of the peripheral blood, given multiple complex barriers between these compartments [10]. CSF cells are primarily of hematopoietic origin. In contrast to the peripheral blood, healthy CSF cellular composition is predominantly lymphoid cells such as CD4+ T cells, whereas myeloid lineage cells account for a lower proportion [11]. Few cells are found in CSF compared to blood, with estimates ranging between 1-3 cells/μL [12] in healthy individuals.

Single-cell-based transcriptomic and epigenetic methods provide high resolution and depth of information about rare or underrepresented cell populations, unbiased assessment of cellular heterogeneity, cell lineage and trajectory, and modeling of intercellular communications. With the advent of single-cell RNA sequencing (scRNAseq), new CSF cell populations have been identified [13,14], and possible mechanisms of neurodegenerative disease states have been elucidated [15,16], with far greater sensitivity and depth of discovery than that enabled by related cellular-based approaches such as flow cytometry [17]. Although using freshly acquired CSF cells may be thought of as ideal, cryopreservation of CSF cells has many practical advantages, such as the opportunity to provide inventory to biorepositories and share resources, reduction of batch effects in cross-sectional analyses, and providing a means to perform longitudinal intra-individual comparisons. However, there are concerns that the transcriptional makeup of the cells will change during the cryopreservation or the thawing process. While there are reagents available that could simplify and reduce batch effects in large studies through sample fixation at the time of collection (e.g., 10x Genomics Fixed RNA Profiling [18]), such approaches rely on probing against a limited subset of genetic targets [19] and may prohibit clonal analyses. To date, several non-CSF-related studies have compared the quality of scRNAseq transcriptomic results between cryopreserved and freshly captured cells using cells derived from peripheral compartments including blood [20–25], generally showing the success of recapitulating fresh cell-based analyses in cryopreserved samples. However, to our knowledge, only two studies have compared fresh versus cryopreserved CSF cells [26,27]. The first study that demonstrated the feasibility of using cryopreserved CSF scRNAseq did not directly compare fresh vs. cryopreservation [26], whereas, the other study is limited to a technical evaluation with no biological findings [27].

Cryopreservation agents include methanol, laboratory-prepared serum combined with dimethyl sulfoxide (DMSO), or a purchasable storage medium (e.g., Recovery). Epigenetic analysis can benefit from additional sample preparation, such as DNase incubation [28]. Here, we compared transcriptomics results from multiple CSF donors seen in the outpatient setting to evaluate possible normal pressure hydrocephalus (NPH). Their evaluation includes a large volume lumbar puncture. Owing to the excess of fluid collected, we could divide CSF from each donor sample into a fresh and one or more cryopreservation method samples, enabling a thorough comparison of fresh to cryopreserved CSF cells. We report on cellular quality, cell-type proportions, and differences in overall gene expression patterns by condition. we demonstrate the utility and compatibility of various CSF cryopreservation techniques for identifying latent genetically based differences in cellular transcription patterns.

## Methods

### Clinical methods

CSF samples were collected as excess fluid from lumbar punctures of patients receiving care at the Mass General Hospital Memory Disorders Unit, as part of an NPH-related clinical evaluation. All lumbar punctures were performed by SEA, using a 24-gauge Sprotte spinal needle, with approximately 20-35mL CSF removed. The first several mL were used for clinical labs and avoided for research purposes, given the risk of procedure-associated red blood cell (RBC) contamination. Massachusetts General Brigham Institutional Review Board approved research use, and informed consent was obtained from all donors.

### CSF cell collection and storage

Fluid was collected in sterile 15 mL Falcon tubes and transferred on ice (approx. 30-45 min) prior to centrifugation. Samples were centrifuged at 300xg for 10 min at 4°C using a swinging bucket rotor. All but 0.1 mL of fluid was removed, and the cell pellet was resuspended. Cell viability and cell count were performed using Luna automated cell counter with acridine orange and propidium iodide viability dye. Red blood cell contamination was determined by subtracting the nucleated cells from the total cells. cell suspension was divided accordingly into one of the below cryopreservation conditions alongside the fresh CSF cell fraction, with a minimum of 5000 cells per condition. Fresh (FRE) CSF samples were immediately prepared for running on the 10x controller. For cryopreservation conditions, the divided cellular fraction(s) was cryopreserved at −80C in equal volumes of the following cryoprotectants: 1) FBS - cells were stored in ice-cold Fetal Bovine Serum (FBS, Thermo-scientific, cat# A4766801) mixed with 10% DMSO (cat# D2650), 2) REC – cells were stored in Recovery media (Thermo-scientific, Cat# 12648010), 3) DNA – Cells were treated with 1X DNase (Stemcell, cat# 07900) for 30 minutes at 22°C, then washed and spun at 3000xg for 6 minutes, followed by cryopreservation in FBS mixed with 10% DMSO. Use of DNase is commonly recommended for some epigenetic analyses [28]. Although we do not examine epigenetics here, we provide quality control data regarding the use of DNase before FBS/DMSO cryopreservation. Methanol cryopreservation based on previously established 10x protocols [29] led to poor thawed cell recovery, leading us not to pursue this method further. Cryopreserved cells were warmed to near complete thaw in a 37°C water bath, after which pre-warmed wash buffer (phosphate-buffered saline supplemented with 1% bovine serum albumin and RNase inhibitor) was added (1.5 mL) to the cryovial. Cells were resuspended and transferred to a 15mL falcon tube containing 9mL warmed wash buffer. The cryovial was rinsed again with warmed wash buffer for further cell recovery). Falcon tubes were centrifuged at 300xg at 4°C for 5 min. The supernatant was removed, and cells were resuspended in 500 μL wash buffer and transferred to a smaller 1.5 ml tube; 30 μL of the aliquot was removed and analyzed for viability and cell counts using Calcein-AM and NucBlue Live ReadyProbes Reagent (Invitrogen, Carlsbad, CA). The sample was centrifuged at 300xg at 4°C for 5 min, resuspended in wash buffer, and loaded into the 10x 5’ v2 mix sans reverse transcriptase enzyme. Finally, reverse transcriptase was added, and the sample was loaded onto the 10x chip K. See Fig 1a for CSF workflow. The list of subjects, as well as the type of cryopreservation method(s) performed on each subject’s CSF cells are listed in S1 Table. The number of tested cryopreservation conditions was determined based on the total cell count. Cryopreserved samples were prepared in parallel with the fresh running on the 10X controller and run later after thawing and counting, within 12 months of initial cryopreservation. Fresh or thawed cryopreserved CSF mixed with 10x 5’ v2 RT Reagent Master Mix were immediately processed with the 10x Genomics Chromium Next GEM Single Cell 5’ v2 kit (10x Genomics, Pleasanton, CA), cDNA amplified, and library constructed per the manufacturer’s protocol. Library quality control (QC) was based on Agilent Tapestation 4200 HS D1000 screentapes (Agilent Technologies, Waldbronn, Germany). Multiplexed library pool QC was based on Agilent Tapestation 4200 HS D1000 and Kapa Library Quantification Kit for Illumina platforms (Kapa BioSystems, Boston, MA) and sequenced at shallow depths on Illumina’s iSeq 100 v2 flow cell for 26×10×10×90 cycles for estimated reads per cell. After demultiplexing, libraries were rebalanced based on reads per cell. Normalized pool QC was based on Agilent Tapestation 4200 HS D1000 and Kapa Library Quantification Kit for Illumina platforms and high depth sequenced on Illumina’s NovaSeq 6000 S4 v1.5 flow cell for 26×10×10×90 cycles, to obtain a sequencing depth of 50K reads/cell (whole transcriptome libraries). Reads were aligned to reference 2020-A (Ensembl 98) transcriptome and quantified using Cell Ranger v6.0.2 multi pipeline. Further data processing and analysis were performed using the Seurat library (v 4.3.0) [30–33]. ggplot2 [34] package was used to generate the visualizations in conjunction with Seurat package. Quality control was performed on each sample: cells with less than 200 genes, more than 4500 genes (presumed doublets), or greater than 20% mitochondrial transcripts were filtered from further analysis. The average number of genes per cell was 1578. Red Blood Cell (RBC) (defined as the cells that showed high expression of Hemoglobin genes HBA and HBB) were manually removed from the analysis.

**Fig 1.**
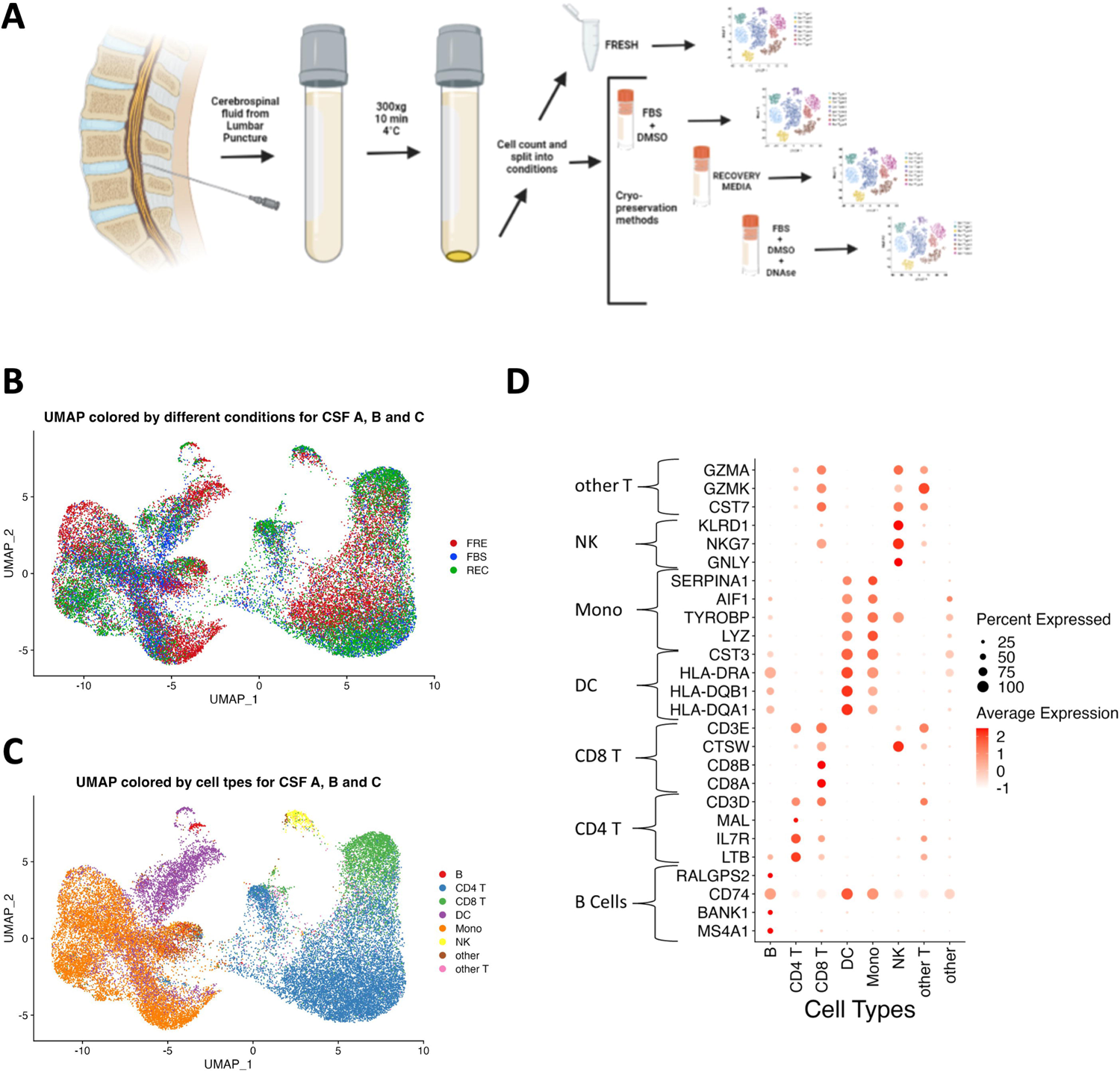
scRNAseq of fresh and cryopreserved CSF cells. a) Graphical schema depicting the experimental design. Freshly collected CSF was spun and the resulting CSF cells were counted, cells were then immediately divided into either FRE which was immediately processed on the 10x Chromium Controller to capture single cells or cryopreserved with protocols for FBS and/or REC and/or DNA for downstream processing basing on the total cell count. The cryopreserved fractions were later thawed and run on the 10x Chromium Controller, and the resulting data were analyzed together. b) Uniform Manifold Approximation and Projection (UMAP) plot of all the cells of samples from CSF A, B, and C, annotated to depict the FRE or FBS or REC protocols. c) UMAP plot of all the cells of samples from CSF A, B, and C in protocols FRE, FBS and REC annotated to depict the cell types clustered using Azimuth peripheral blood mononuclear cells (PBMC) reference atlas at L1 resolution. d) Dot plot depicting the concordance between respective cell-type specific enriched marker gene expression of all the cells from samples CSF A, B, and C (on the y-axis) as characterized by earlier published CSF scRNAseq studies that are correspondingly identified and clustered accordingly (on the x-axis) by Azimuth reference atlas at L1 resolution.

### Cell integration and data processing

Samples were integrated using the Seurat package v4.0 [35]. First *FindVariableFeatures()* was used to find the top 2000 highly variable genes. Then *SelectIntegrationFeatures()* was used to select features and PCA was performed using *RunPCA()* using the selected features. Reciprocal PCA was performed to identify pairwise anchors between the reference and the list of datasets using *FindIntegrationAnchors()*. Two fresh samples (CSF A and CSF C), one male and one female, both with high total counts of cells, were selected as anchors for data integration. Integration was performed using *IntegrateData(). NormalizeData()* was used for normalizing the count data for each dataset, with a scaling factor = 10000. Data was log-transformed before analysis. We used Azimuth [31], a reference atlas of human peripheral blood mononuclear cells (PBMCs), to define and annotate the CSF cell types. *RunAzimuth()* was used to map our data to PBMC reference dataset. This returned a Seurat object that contained cell annotations (at multiple levels of resolution), and prediction (i.e., confidence) scores for each annotation: This score was used for filtering cells with low confidence classifications. To validate the results of Azimuth-based cell type calling, we compared the Azimuth cell type annotations with commonly used immune cell marker genes, taken from recent CSF-based studies [15,16,36].

Normalized gene expression for these markers was further plotted using *Dotplot()* from the ggplot2 package. For obtaining overall gene expression correlation plots, average gene expression was calculated for each subject (CSF A, CSF B and CSF C) by each condition. Pearson method of correlation was used and the plots were visualized using *ggscatter()* in the ggplot2 package in R. Euclidean distance matrix was computed using *dist()* in stats (version 4.2.1). Hierarchical clustering was performed on the distance matrix by the complete clustering method using *hclust()* in the stats package. We picked the genes HSPA1A, HSPA1B and HSP90AA1, which are related to cell stress response pathways (GO:0031072 - heat shock protein binding and GO:0051082 - unfolded protein binding) as published in an earlier study [24], to compare the effect of cryopreservation. Global gene expression was plotted using Seurat *VlnPlot()*. Differential gene expression analysis was performed between two conditions within a cell cluster using Seurat *FindMarkers().* Significance was defined as Bonferroni-adjusted p-value <0.05 and absolute log2 fold change >0.58 (fold change >1.5). We further picked the genes XIST, KDM4D, UTY, DDX3Y and USP9Y, related to sex differences as published in the earlier study [37], to find the similarities between differential expression analysis among all the conditions together, as well as in each individual condition. *pheatmap()* was used to plot the heatmap of differential expression.

### Statistical analysis

Statistical tests were indicated where applicable for comparing quality control metrics. One-way ANOVA and Tukey’s multiple comparisons test were used for normally distributed data, and Friedman’s was used for the comparison of percentages. Statistical tests were performed with GraphPad Prism v9.5.

## RESULTS

### Donor demographic and clinical characteristics

Donor ages ranged from 61-85 years (median age 74 years) at CSF collection. According to self-report data recorded in electronic health records, all donors identified themselves as white/Caucasian and 55% of donors were male. See S1 Table for individual donor demographic and clinical information.

### CSF sample characteristics

Average CSF volume was 28.7 mL (SD 8.04, range: 8-38 mL). Total CSF cell counts averaged 1980/mL, (SD 1810, range: 570-6700/mL), RBC contamination was overall low, averaging 754/mL, (SD 105, range: 0-305/mL). These data exclude two samples, one for which RBC contamination in one sample was unusually high, enabling visualization of a red pellet. This sample was removed from further analyses in this study. See S1 Fig for examples of cell pellets from spun CSF samples with and without RBC contamination. The other sample with notably high RBC contamination was CSF F (43,500 RBCs/mL of CSF). We included this sample for limited analysis (FBS vs REC comparison). CSF cell viability averaged 99.29%, (SD 1.20, range 96.75-100%). Cells per condition averaged 10,600, (SD 2800, range: 5000-16000 cells). Samples showing pleocytosis were divided into multiple cryopreservation conditions. Sample details by donor and cryopreservation method can be found in S1 Table. A total of 7 FRE, 11 FBS, 11 REC and 5 DNA, were obtained. Donors A, B, and C each contributed at least one sample to the FRE, FBS, and REC conditions, allowing for a within-subjects comparison of these conditions. Donors A, B, C, and J contributed samples for FRE vs FBS comparison. Donors A, B, C, and H contributed samples for FRE vs REC comparison. Donors A, B, C, D, E, F, and G contributed samples for FBS vs REC comparison. Donors A, C, D, F, and G contributed samples for FBS vs DNA comparison.

### Single-cell sample quality control metrics

Pre-filtering, total droplet-encapsulated cell count was 78,551, summing 16,279 FRE, 24,850 FBS, 27,756 REC, and 9666 DNA. Post-filtering by gene and mitochondrial transcript count led to an overall cell count reduction of 1%. 21,117 genes were detected on average across all samples, with 19,173 in FRE, 19,428 in FBS, 19,747 in REC, and 17,416 in DNA. The pre-filtration average unique molecular identifiers (UMI) were 4,576 across all samples, with 6,230 in FRE, 4,062 in FBS, 3,788 in REC, and 5,248 in DNA. Filtration led to an increase in overall average UMI by 1%. Pre-filtered average percent mito-based reads were 2.6% overall, with 1.9% in FRE, 2.4% in FBS, 3.5% in REC, and 2.6% in DNAse. Filtration led to a decrease in average percent mitochondrial reads of 1%. No significant differences were found between treatment conditions, or in comparing pre-vs post-filtration metrics. We focus on comparisons between CSF A, B, and C in our studies since these were the only donors for which the FRE, FBS, and REC conditions were all available. CSF A, B, and C feature count (average genes per cell), UMI count, and percent mitochondrial read distributions are graphed in S2a-c Figs. The average genes detected per cell in 2041 in FRE, 1102 in FBS and 1273 in REC. There was an overall decrease in the average genes detected per cell in both the cryopreservation conditions, but REC was found to be significant (one way ANOVA, Tukey’s Posthoc test, p =0.0031). UMI was overall higher in FRE than FBS and REC groups, though this was not significant statistically: F(1.20, 2.401) = 8.86, p=0.08. There were no differences in the percentage of mitochondrial reads across the three groups (p=0.94). Features, UMI, and percent mitochondrial reads were very similar between FBS and REC (including samples CSF A-G, see S4b Fig), and between FBS and DNA conditions (including samples CSF A, C, D, and G, see S5b Fig). See S2 Table for detailed quality control-based metrics for each donor, condition, and sample replicate. Examination of global heat shock protein genes HSPA1A, HSPA1B, and HSP90AA1 found minimal to no differences in gene expression by cryopreservation method in any of the three donors (S2d Fig).

### CSF cell types

Integrated CSF cells from donors A, B, and C, colored by fresh or cryopreservation conditions, are visualized by UMAP plot in Fig 1B. The cell types defined by Azimuth L1 supervised clustering are shown in Fig 1C. The samples represent all L1 cell types, including CD4 and CD8 T cells, monocytes, dendritic cells, NK cells, and scant B-cells. Few cells fell into ‘other’ or ‘other-T’ categories by the L1 algorithm. We provide several methods of validation of the Azimuth-based method. First, we cross-validated Azimuth-based cell designations to cell type markers from the literature (see METHODS). The Azimuth L1-based cell type predictions are concordant with established phenotypic markers. Next, we prepared an adjacency table by donor, condition, sample, and Azimuth-derived cell type. L1- and L2-resolution cell designations are provided (L2 information is provided as an additional resource but is not utilized for further data analysis in this study). Mean predicted scores, a confidence metric in cell type callings by Azimuth, ranged from 78-88%. By cell type (%) in descending order of percent presence in sample: CD4 T (87), Mono (83), CD8 T (87), DC (11), B (78), Other (57), Other T (67), and NK (78). The cryopreservation method (including FBS, REC, DNA) did not affect mean predicted scores (S3 Table).

### Effects of cryopreservation method on cellular proportions

UMAP and stacked bar-plots of the Azimuth L1-based cell type by cryopreservation method and donor are shown in Fig 2A and Fig 2B respectively. Overall, all cell types were identified across all donors and conditions. There were generally even distributions of cell proportions by cryopreservation method across donors. Based on the comparison of cell type proportions (Fig 2b) FBS cryopreserved samples appeared to have slightly greater proportions of monocytes and lower proportions of CD4 T cells than FRE or REC samples across donors. However, this was not found to be significant by statistical test (monocytes: p=0.19, and CD4 T cells: p=0.19). Examining the raw cell counts (S2 Table), these proportional differences seem to be related to a selective loss of lymphocytic cells. For instance, in two independently prepared samples of CSF A FRE, there were 1266 and 1240 monocytes, with 1233 and 1308 CD4 cells, respectively. The FBS samples had fewer overall cells, with 1059 and 1888 monocytes, but fewer CD4 cells: 958 and 833, as compared to monocytes. DNAse-treated samples, cryopreserved similarly to FBS samples, showed fewer CD4 cells: 866 and 1190 than monocytes: 1164 and 1471. REC cryopreservation looked more like FRE: 1119 and 780 monocytes, with 1089 and 1151 CD4 cells. Comparing FBS to REC (S4c Fig), the phenomenon of slightly lower CD4 T cell proportions in FBS than REC is maintained. There is no consistent difference between cell proportions in comparing DNase pre-treated samples to FBS (S5c Fig).

**Fig 2.**
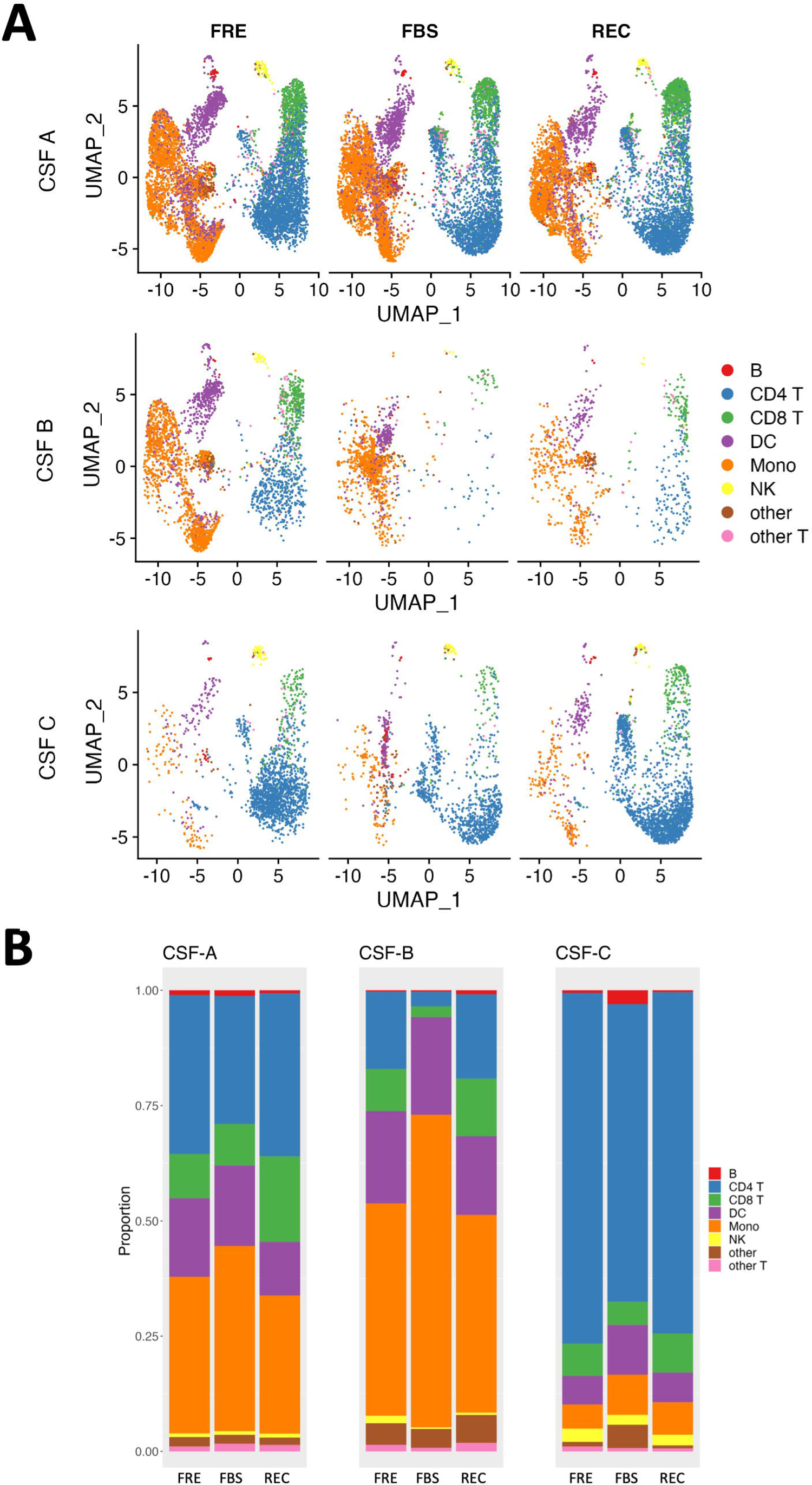
Distribution and cellular type proportions of cryopreserved versus freshly captured CSF cells. a) UMAP plot of all the cells in samples CSF A, B, and C from FRE, FBS, and REC protocols were annotated to show the cell types clustered using Azimuth PBMC reference atlas [35] at the L1 resolution. b) Stacked bar plots depicting the individual cell type proportions obtained after clustering using Azimuth reference atlas at the L1 resolution in samples CSF A, B and C from FRE, FBS and REC protocols.

In contrast to the limited differences in cell proportions by cryopreservation method, donor-specific biological differences are preserved, and marked. For instance, CSF C shows far greater proportion of CD4 T cells, and proportionately fewer monocytes than CSF A and B. A dotplot stratifying CSF samples by cryopreservation method and Azimuth L1 cell type is presented in S3 Fig. Dots showing average gene expression reveal highly similar cell type marker expression and percent expression across cryopreservation methods compared to the fresh condition.

### Effects of cryopreservation method on global gene expression

To answer whether the cryopreservation method affected global gene expression, we prepared a hierarchical clustering dendrogram comparing global (across all cell types) gene expression across FRE, FBS, and REC conditions for CSF A, B, and C (Fig 3a). Samples obtained from the same donor using different cryopreservation methods clustered together, indicating that the observed variations in global gene expression profiles are primarily due to biological variability rather than technical factors (similarity matrix is provided in S4 Table, lower scores indicate closer cluster distance). We next correlated global gene expression within donors between FRE and cryopreserved conditions (Fig 3b), finding very high correlations between FRE and FBS, and FRE and REC conditions (average R-value: 0.98, Range: 0.94 – 1).

**Fig 3.**
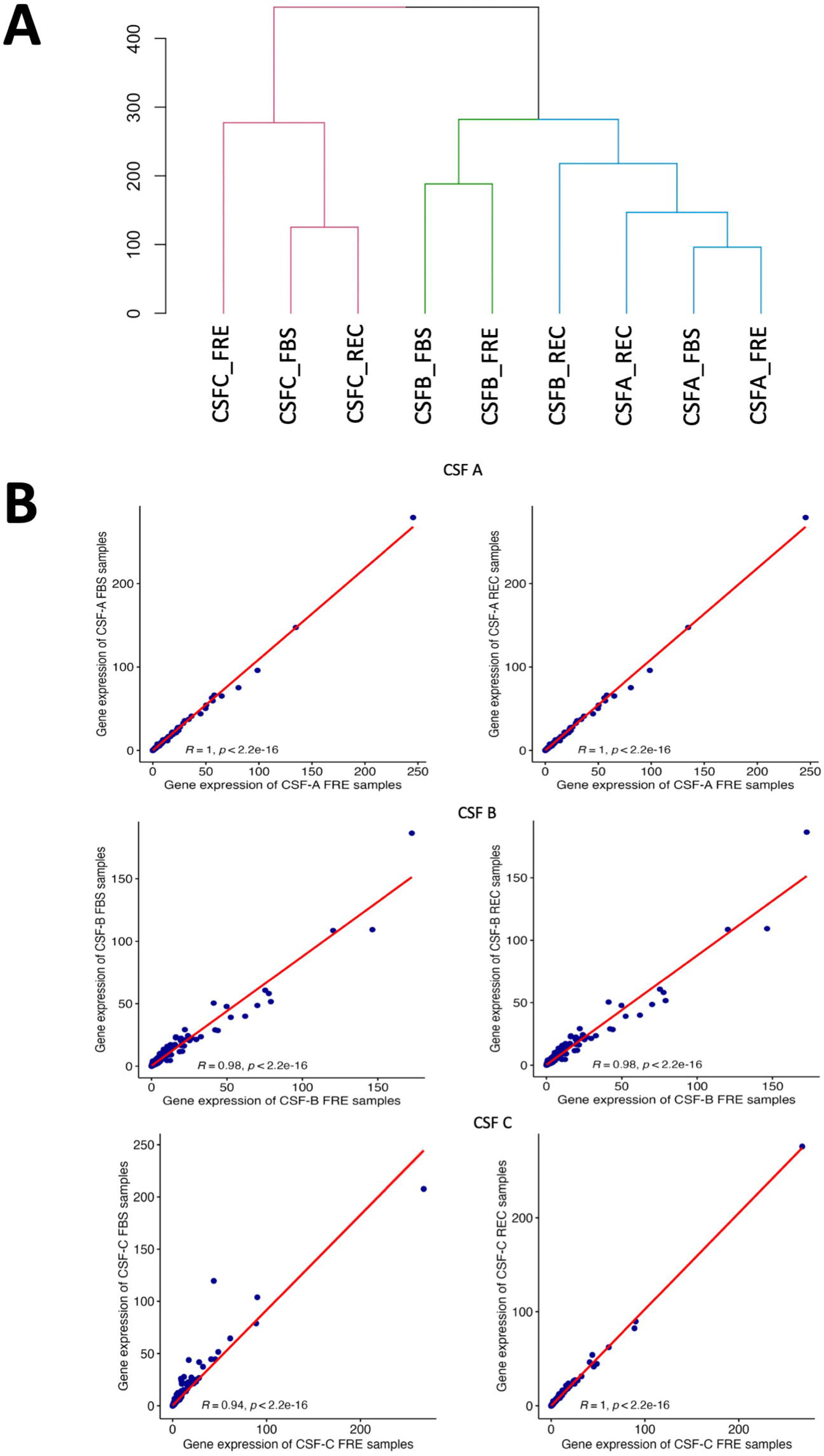
Comparison of overall gene expression between cryopreserved and fresh CSF samples. a). Dendrogram showing the hierarchical clustering of relative sample to sample distances in samples CSF A, B, and C in FRE, FBS, and REC protocols. Note the companion adjacency matrix found in S4 Table. b). Scatter plots with correlation statistics of FRE vs FBS and FRE vs REC global gene expression in samples CSF A, B and C.

### Effect of cryopreservation on revealing latent sex-linked expression

To test whether the cryopreservation method impacted our ability to extrapolate functional or biologically meaningful data, we utilized established knowledge regarding sexually dimorphic gene expression: XIST found on the X-chromosome and KDM4D, UTY, DDX3Y, and USP9Y on the Y chromosome. We analyzed CSF A, B, and C samples (FRE, FBS, and REC conditions) to identify differentially expressed genes between males and females (Fig 4A). We evaluated CD4 cells as exemplars given their abundance in CSF. We observe many sexually dimorphic genes (n = 149 upregulated and 124 downregulated genes based on male/female fold change). Each sex-specific gene of interest was in the expected direction (genes shown in red on the right side of the volcano plot indicate higher expression in males). These results demonstrate that combining samples prepared using different cryopreservation strategies did not affect the detection of sexually dimorphic gene expression. We next determined whether cryopreservation-specific conditions are independently capable of detecting sexually dimorphic gene expression differences. To do this, we collapsed all cell types together and compared cryopreservation conditions. As seen in Fig 4b, the fold change differences between sexes for the five sex-linked genes of interest showed highly similar patterns across the various cryopreservation methods. They compared favorably to combinatory and FRE results.

**Fig 4.**
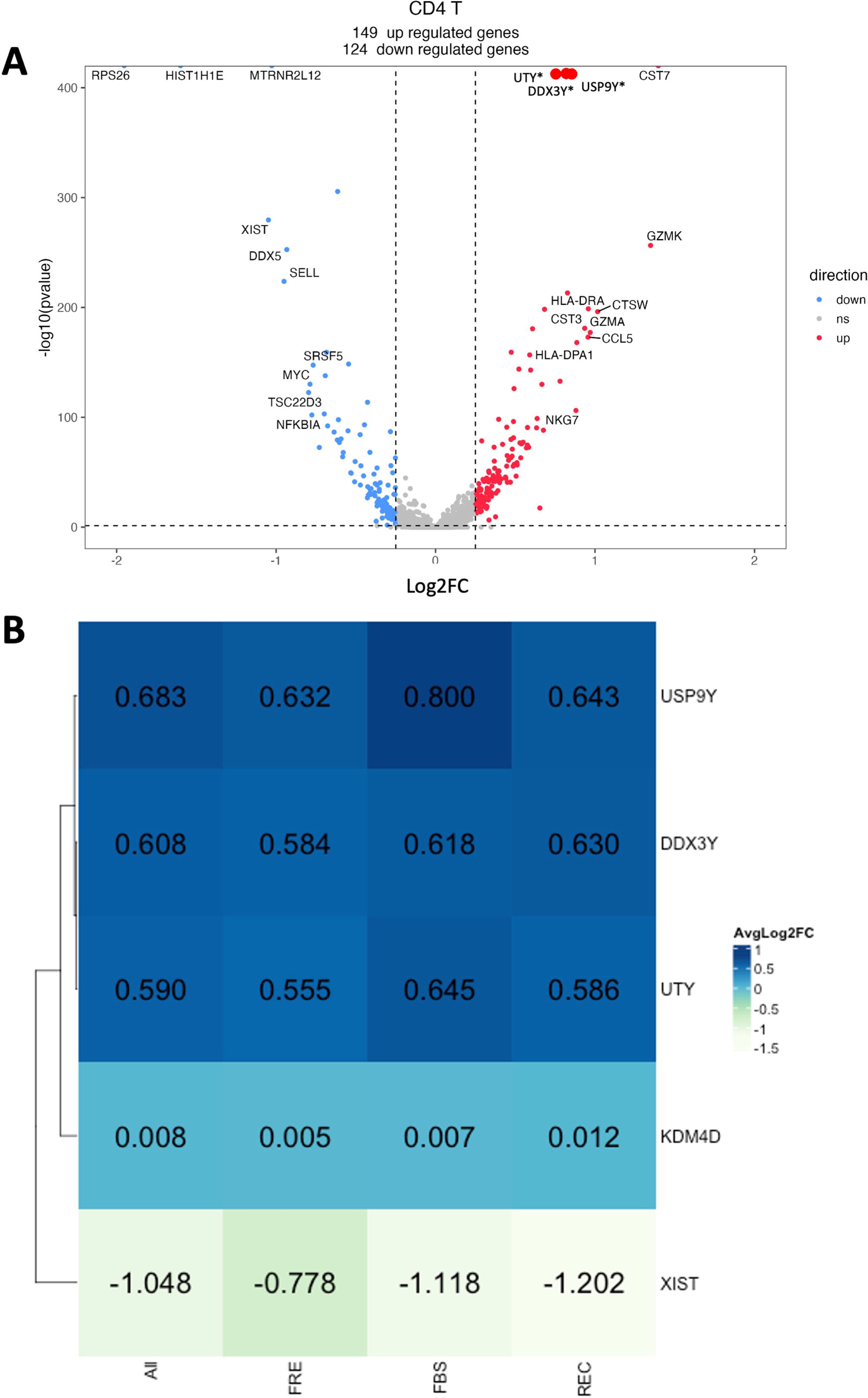
Differential gene expression analysis of sexually dimorphic genes from fresh and cryopreserved CSF samples. a) Volcano plot showing the significant (p-adjusted < 0.05, Log2FC > 0.58) differentially expressed genes between males and females in CSF A, B, and C CD4+ T-cells combining all fresh and frozen samples. Genes upregulated in males are colored red, and downregulated in males are colored blue. Grey dots indicate the non-significant genes. b) Heatmap showing the average log2fold changes of the sexually dimorphic genes XIST, KDM4D, UTY, DDX3Y, and USP9Y from differential gene expression analysis of CD4+ T-cells in CSF A, B and C for All samples (FRE, FBS and REC combined), and stratified by FRE, FBS, and REC.

## Discussion

In this methodological study of CSF cells donated by patients receiving evaluations for possible normal pressure hydrocephalus, we compared various cryopreservation approaches to freshly run CSF cells through the scRNAseq process. We used three cryopreservation methods, including two cryopreservation agents (FBS/DMSO and Recovery media). We included an additional DNase step prior to FBS/DMSO cryopreservation given this step’s known benefit on the quality of epigenetic analyses. Overall, we found little impact of cryopreservation on cell integrity and stress, cell type representation, or overall gene expression. Nuanced findings of the study include possible relative losses of CD4+ T-cells using the FBS/DMSO cryopreservation technique, and limited loss of sequencing depth, as seen by relatively lower average UMI in all cryopreserved samples. The robustness and interchangeability of cryopreserved and freshly run samples was demonstrated using an example of sexually dimorphic gene expression, which was noted using a combined dataset from fresh and pre-frozen samples, as well as by stratified analysis by cryopreservation method. When compared to the fresh condition, there were no apparent differences due to cryopreservation methods using FBS/DMSO, Recovery media, or from prior DNase treatment. Our data support the ability to cryopreserve and later utilize CSF cell samples, enabling batched single-cell analyses for wide-ranging cellular and molecular-based advances in neurological, oncological, infectious disease, and psychiatric research.

### Effect of cryopreservation on cellular stress, and sequencing quality

In the present study, we noted limited to no impacts of cryopreservation on evidence of cell stress, and our data supported overall robust single cell transcriptome results. For comparison, in the preprinted study by Touil et al., [27], researchers recovered approximately 70% of cryopreserved cells, without other notable impacts of cryopreservation on mitochondrial reads. We similarly routinely find approximately 70% cell recovery (not shown) post-thaw, and likewise do not find any evidence of increased cellular stress, as represented by the proportion of mitochondrial reads. We also find that global expression of cellular stress response proteins, including heat-shock proteins HSPA1A, HSPA1B, and HSP90AA1 was unaffected by FBS or REC cryopreservation. Previous studies have noted upregulation of cellular stress pathways in cryopreserved dissociated tumors [24] or single-cell subpopulations of peripheral blood mononuclear cells. PBMCs [22]. The latter study noted that while the effects appeared limited and did not disrupt key aspects of the cellular subpopulation, cell populations showing evidence of cellular stress clustered uniquely from healthy cells, allowing for selective filtration. Differences in thawing and cryopreservation strategies used across ours and previous studies are likely to account for differences in cellular stress pathway activation; this supports the importance of consistency in sample treatment surrounding cryopreservation practice (e.g., temperature control, spin speed, and freezing and thawing schedule).

Non-viable cells are readily identified using hemocytometers and dye markers. Generally, we find >90% cell viability in CSF cells. The differences between recovery and viability are likely related to the degradation of damaged cells during post-thaw washes. The report by Touil et al., [27] also notes higher hemoglobin gene expression in fresh samples despite excluding RBCs from analyses. Our dataset reveals a similar phenomenon (not shown). We further note from other optimization studies in our labs that increasing the temperature of centrifugation (i.e., room temperature) decreases overall cell viability substantially (20-30% pre-freeze), with the greatest effect on RBC viability (data not shown). This selective effect can mask peripheral blood contamination, leading to a false interpretation of peripheral contamination as CSF-specific cells. Use of an atraumatic LP technique and avoiding using the first several mL of collected CSF can prevent peripheral blood contamination, and maintenance of 4C of the sample through cryopreservation is essential. Unbiased identification and filtration of cells containing unexpected or ambient gene expression, such as hemoglobin, may be practically useful, but should also be applied with caution given the underlying possibility of such findings reflecting peripheral blood contamination. Finally, both groups identified a limited but consistent decrease in UMIs and genes from cryopreservation. We find this to be consistent between FBS/DMSO and Recovery media as compared to Fresh samples. We attribute this to slight RNA degradation occurring in the lengthier processing and sample preparation time required during thawed samples.

### Effects of cryopreservation on cellular proportion

We find very limited effects of cryopreservation on cell proportions. A careful evaluation revealed non-statistically significant relative losses of CD4+ T cells in FBS/DMSO cryopreserved samples versus fresh samples, with sparing of this phenomenon using Recovery media. Oh et al., [26] compared Recovery to fresh, and did not evaluate FBS/DMSO. Like our findings, they did not note any impact of Recovery media on CD4+ T cells. Rather, they noted an overall low proportion of myeloid cells (6%) and noted mild losses of myeloid cells post-thaw from Recovery cryopreservation. Our study finds greater proportions of CSF myeloid cells (including an average of 27% monocytes). Several donors had far lower monocyte proportions present in their CSF (e.g., CSF F [3%], and CSF C [6%]). Our findings are unlikely due to miscalling of cell type, given both the substantial variability of monocyte proportions found in our donor population, as well as the high mean predicted score (0.87) of Azimuth nearest-neighbor calling, and our added validation of the Azimuth [31] functions using cell marker genes used for cluster labeling in many scRNAseq studies.

### Limitations

This study has several limitations. First, we recognize that to perform comparative analyses of freshly run versus cryopreserved CSF samples, many cells per donor are required. This may bias our samples towards individuals showing CSF pleocytosis, limiting the applicability of our results to healthy individuals. Next, although we carefully validate our cell populations, a higher standard of cell marker-based identification would rely on a transcriptionally-independent means of cell identification, such as through simultaneous epitope and transcriptome measurement in single cells (i.e., CITE-seq) [27]. Lastly, while we find little impact of cryopreservation on basal CSF cell transcriptomics, we cannot account for differences in cellular function that would readily be revealed by functional cellular assays [32].

## Conclusion

In this methodological study of scRNAseq of CSF cells, we find good comparability between cryopreserved and fresh preparations. Sufficient starting volumes of samples, delicate cell isolation, rigid adherence to cryopreservation and thawing steps, and routine quality control steps allow for the robust study of CSF cells, allowing for reduced assay-based variability, and perhaps allowing for great leaps in diagnostic and mechanistic advances in many research domains relating to the central nervous system’s immune milieu.

## Acknowledgements

Sponsors were not involved in the design and conduct of the study, collection, management, analysis, and interpretation of the data; preparation, review, or approval of the manuscript; or decision to submit the manuscript for publication.

## Supporting information

**S1 Fig. Representative images of the CSF cell pellet visible post-centrifugation.** See TIFF file. A sample devoid of RBC contamination is seen on the left, and one with substantial RBC contamination is seen on the right.

**S2 Fig. Cellular features, UMI, mitochondrial reads, and cellular stress-related gene expression.** See TIFF file. All graphs include data from CSF A, B, and C. (a-c) Density plot showing the distribution of the number of transcriptional features (genes), number of Unique Molecular Identifiers (UMI), and percentage of overall gene expression attributed to mitochondrial genes, respectively. (d-f) Violin plots of normalized gene expression levels of HSPA1A, HSPA1B, and HSP90AA1, respectively.

**S3 Fig. Concordance of cell type markers between cryopreserved and freshly run samples**. See TIFF file. Dot plot comparison of cell-type specific enrichment of marker gene expression in FRE, FBS, and REC protocols for samples CSF A, B, and C. Cell-type specific marker gene expression and associated cell types are listed on the y-axis, and Azimuth reference atlas cell designations at L1 resolution are listed on the x-axis.

**S4 Fig. FBS versus Recovery comparisons.** See TIFF file. (a) UMAP plot of all the cells in samples CSF A, B, C, D, E, F, and G from FBS and REC protocols were annotated to show the cell types that were clustered using Azimuth PBMC reference atlas at L1 resolution. (b) Density plot showing the similar distribution of the number of genes, number of unique transcripts, and percentage of mitochondrial genes in all the cells from FBS and REC protocols in samples CSF A, B C, D, E, F, and G. (c) Stacked bar plot depicting the individual cell type proportions obtained after clustering using Azimuth reference atlas at L1 resolution in samples CSF A, B C, D, E, F and G from FBS and REC protocols.

**S5 Fig. FBS versus FBS with Dnase (DNA) comparisons.** See TIFF file. (a) UMAP plot of all the cells in samples CSF A, C, D, and G from FBS and DNA protocols were annotated to show the cell types clustered using Azimuth PBMC reference atlas at L1 resolution. (b) Density plot showing the similar distribution of the number of genes, number of unique transcripts, and percentage of mitochondrial genes in all the cells from FBS and DNA protocols in samples CSF A, B C, D, E, F, and G. (c) Stacked bar plot depicting the individual cell type proportions obtained after clustering using Azimuth reference atlas at L1 resolution in samples CSF A, C, D, and G from FBS and DNA protocols.

**S1 Table. Donor demographics and sample information.** See xlxs file. Subject IDs, Age, Sex, and number of individual reactions tested from FRE, FBS, REC, and DNA samples listed.

**S2 Table. Quality control metrics pre- and post-filtration.** See xlsx file. QC filtering: Cells having 200 – 4500 umi counts, < 20% mito genes detected were retained. 2000 high variable genes were used for clustering. Columns: Total_cells.before.filtering: cells pre-filtering; cells after filtering: cells remaining after application of above filters; Total_genes: Genes per sample, avg_numi: average number of umi counts before filtering; numi after filtering: average number of umi counts after filtering; avg_mito: average mitochondrial genes in each sample before filtering; mito after filtering: average number of mitochondrial genes in each sample after filtering.

**S3 Table. Cell type designations, proportions and mean predicted scores.** See xlsx file. Mean predicted score is the Azimuth-derived confidence score for a given annotation. ‘All’ refers to the total of all cell types (L1 or L2). Columns E-L reference L1 designations (used throughout the manuscript), and columns M-AP list L2 designations (not used elsewhere in the manuscript). Summary information for samples by fresh or cryopreservation method can be found in the lowest rows.

**S4 Table. Dendrogram adjacency matrix of overall sample gene expression.** See xlsx file. The order of the donors and fresh/cryopreservation condition is based on the unweighted Euclidian distance between samples as visualized on the dendrogram (Fig 3a).

## Notes

### Competing Interest Statement

The authors have declared no competing interest.

### Summary of Updates

Added Supplemental Files

