## Supplementary figures and images for "Cryopreservation of cerebrospinal fluid cells preserves transcriptomics integrity for single-cell analysis"

### Supp Fig 1

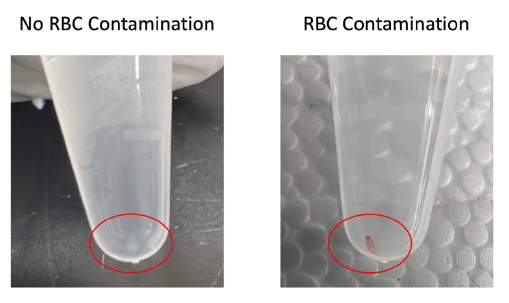

### Supp Fig 2

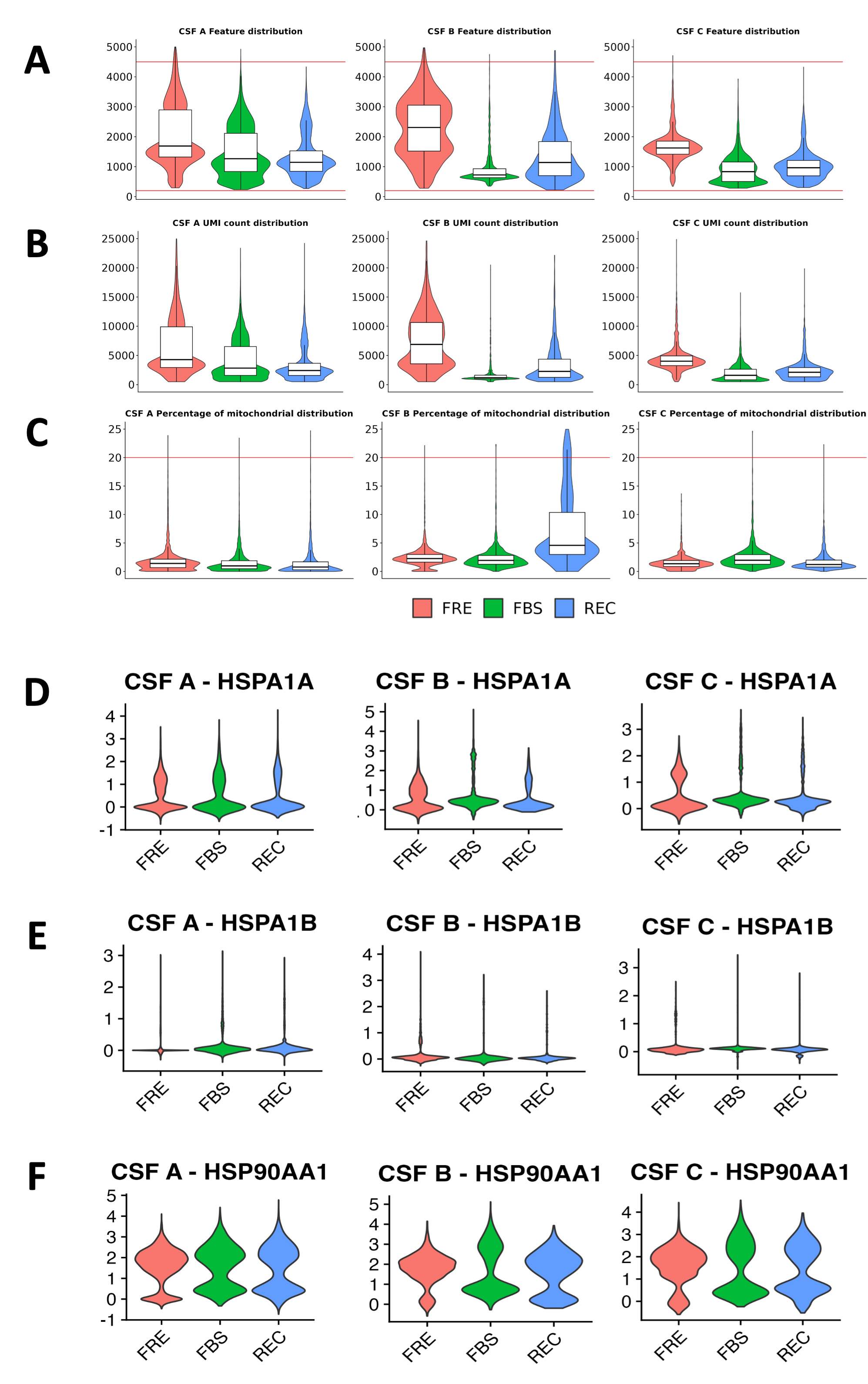

### Supp Fig 3

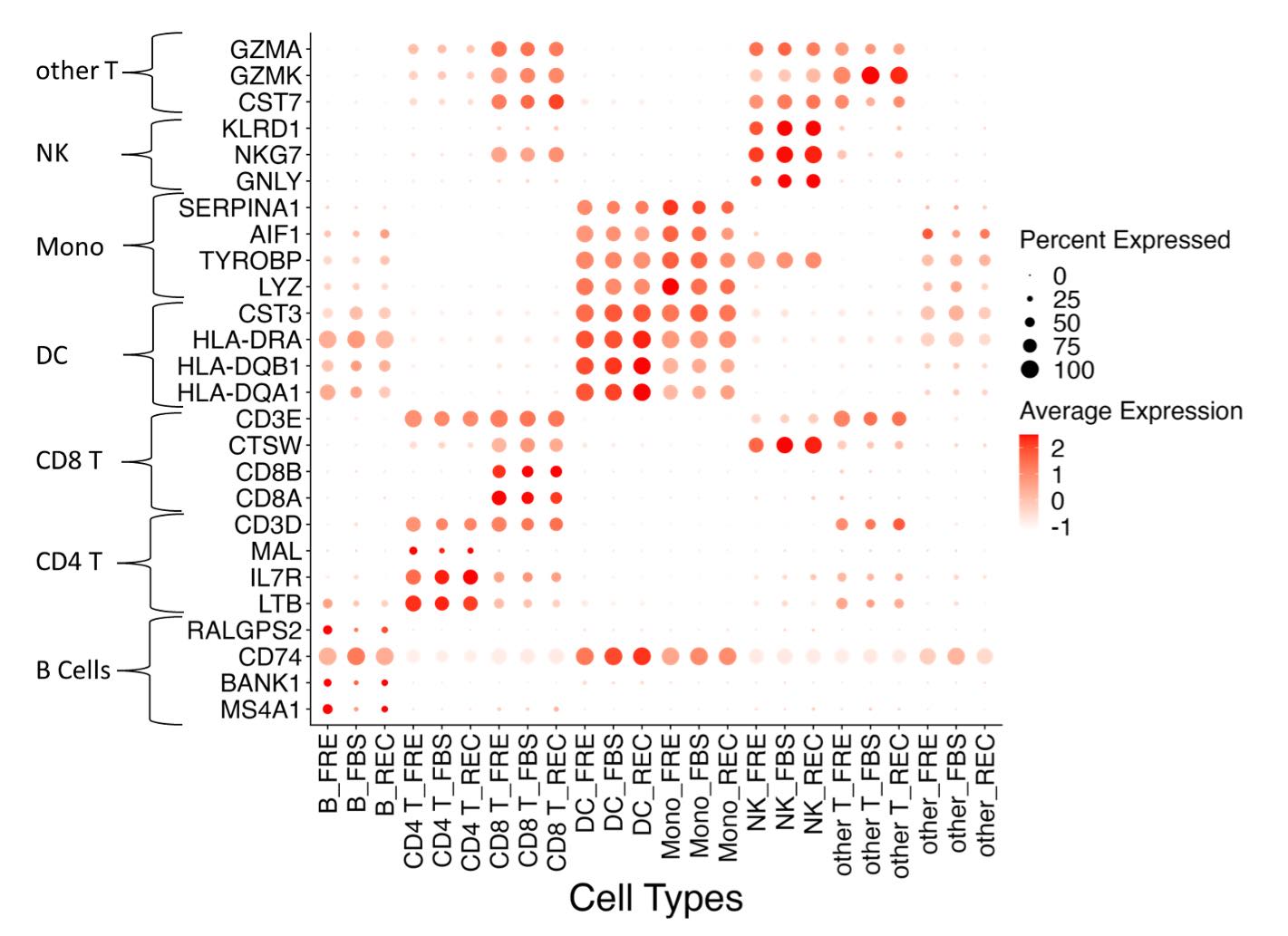

### Supp Fig 4

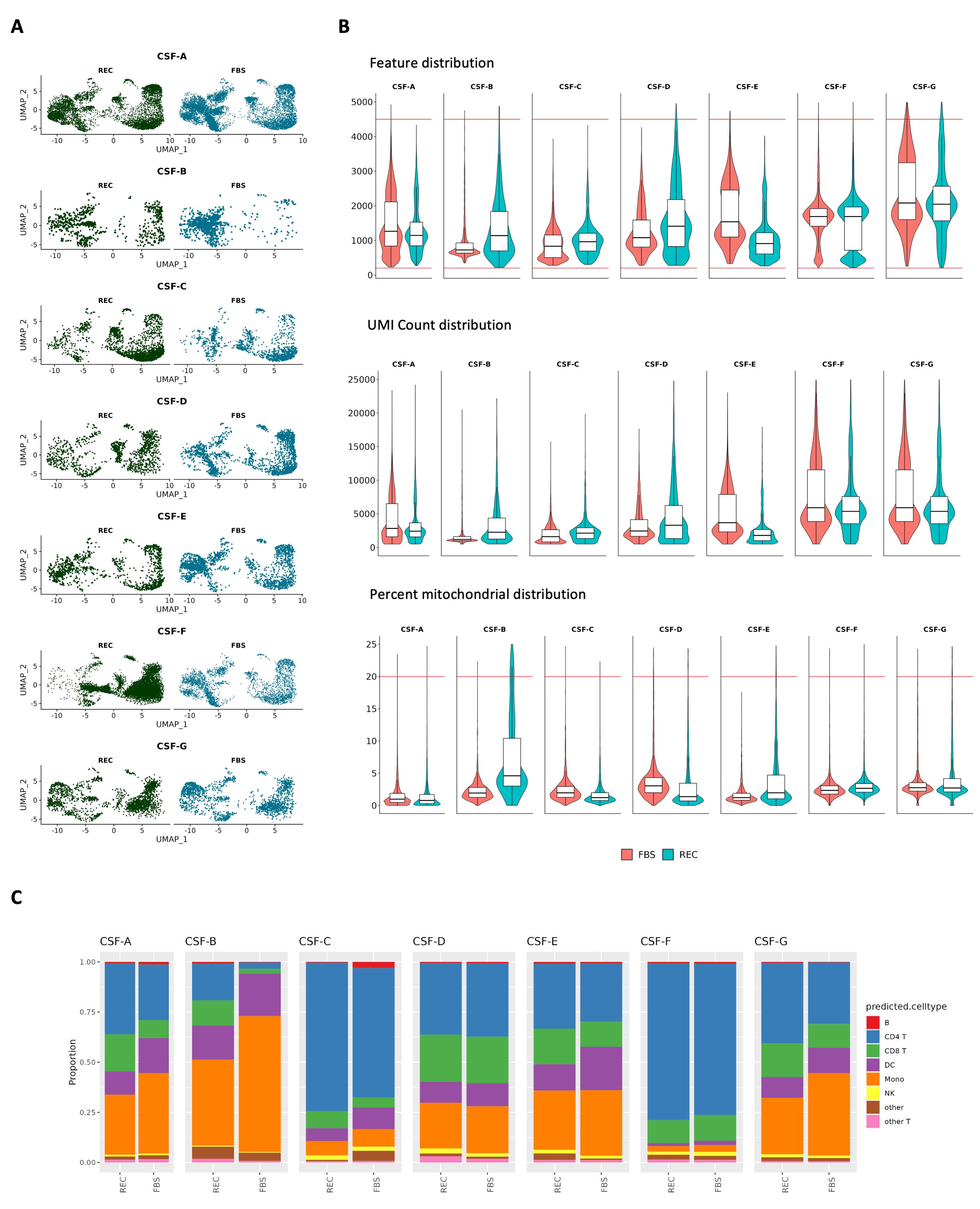

### Supp Fig 5

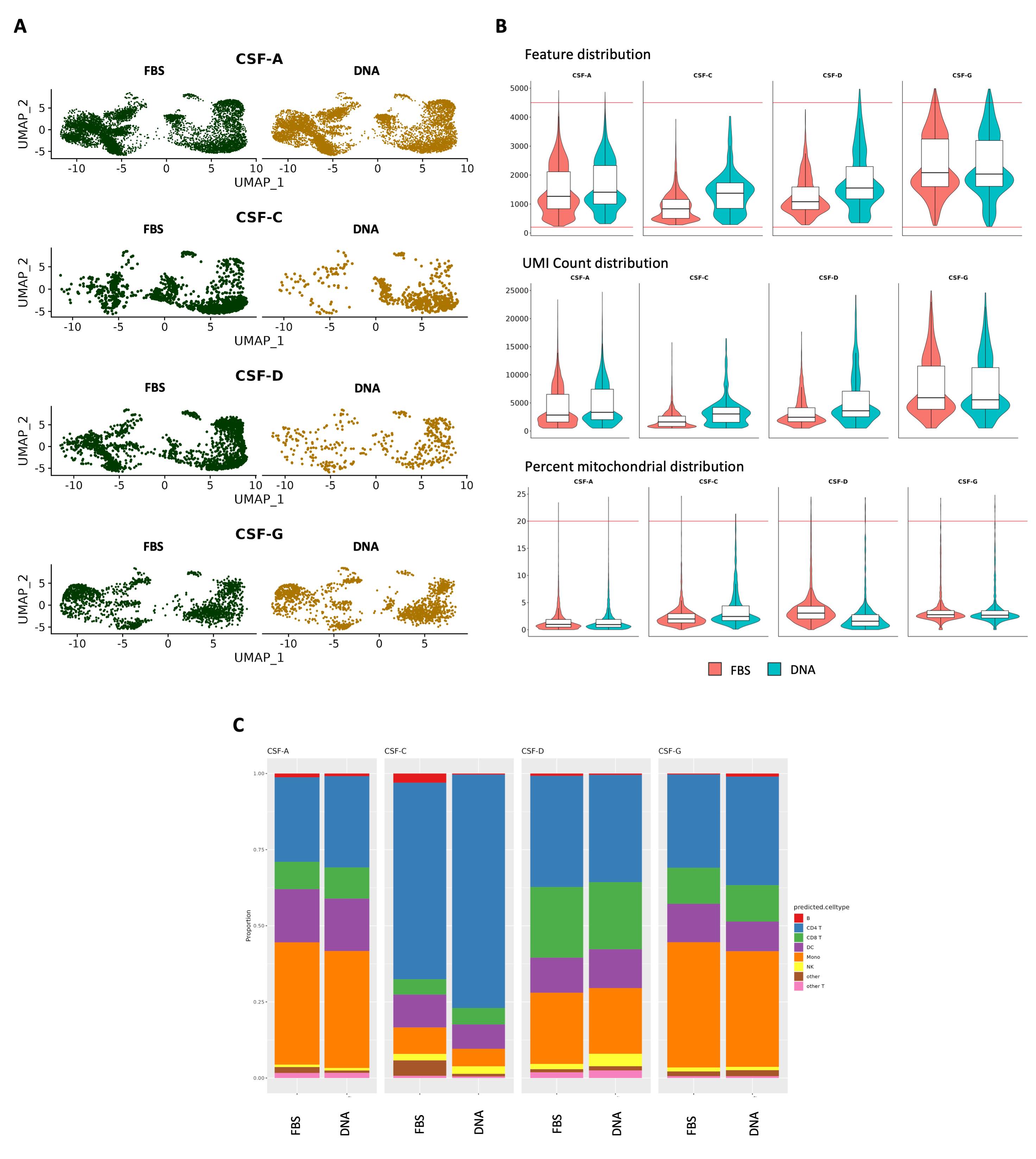
